# High affinity FcγR activating function depends on IRAP^+^ endosomal-signaling platforms

**DOI:** 10.1101/2021.06.24.449774

**Authors:** Samira Benadda, Mathilde Nugues, Marcelle Bens, Mariacristina De Luca, Olivier Pellé, Renato C. Monteiro, Irini Evnouchidou, Loredana Saveanu

**Author notes:** Corresponding author: Loredana SAVEANU, 16 rue Henri Huchard, Faculté de Médecine Xavier Bichat, 75018 Paris, France, **Email:**. Present address: Evox Therapeutics Limited, Oxford Science Park, Robert Robinson Avenue Oxford OX4 4HG (Mariacristina De Luca).

## Abstract

Although endocytosis of cell surface receptors is generally thought to terminate the signaling, for some receptors, endocytosis sustains signaling. We wondered if endosomal signaling participates to the function of the receptors for Fc immunoglobulin fragments (FcRs) that are highly internalized after their activation. We demonstrate here that four different FcRs follow distinct endocytic pathways after activation. While FcαRI is internalized into lysosomes, FcγRIIA is internalized and partially retained in early endosomes, whereas the inhibitory receptor FcγRIIB is internalized in endosomes decorated by the autophagy marker LC3. Interestingly, the high affinity FcγRI is internalized in specialized endosomal compartments described by the Insulin Responsive AminoPeptidase (IRAP), where it remains associated with the active form of the signaling kinase Syk. Our results show that FcγRI has the ability to build endosomal-signaling platforms, which depend on the presence of IRAP and Rab14. Destabilization of the endosomal signaling platforms compromised the ability of peritoneal macrophages to kill tumor cells by antibody-dependent cell mediated cytotoxicity, indicating that FcγRI endosomal signaling is required for the therapeutic action of anti-tumor monoclonal antibodies.

**One sentence summary:** Binding of immune complexes to FcγRI receptor leads to receptor internalization and sustained signaling from endosomes described by the Insulin Responsive AminoPeptidase and the small GTPase Rab14.

## Introduction

Endocytosis modulates the function of cell surface receptors. These receptors detect signals from the environment and trigger signaling cascades, which usually start at the plasma membrane. Frequently, after ligand binding, the ligand-receptor complexes are internalized and this internalization was supposed to be a way to attenuate or terminate the signaling through ligand and receptor degradation. This view changed when it was discovered that endocytosis of receptor tyrosine kinases (RTKs), rather than attenuating the signaling, sustain regulated signal transduction from endosomal compartments (*1*). Later studies showed that many other RTKs and G protein coupled receptors (GPCRs) are able to signal from endosomes, leading to the establishment of the concept of endosomal signaling (*2*).

In the immune system, the concept of endosomal signaling is well documented for endosomal toll-like receptors (TLRs) (*3*). Endosomal TLRs, such as TLR3, 7 and 9, recognize pathogen-derived nucleic acids, but have also the potential to recognize self nucleic acids leading to autoimmune disease. To avoid autoimmunity, their signaling is tightly controlled by several mechanisms. These include steady-state retention of the receptors in the endoplasmic reticulum and their transport to endosomes only upon cell activation. Furthermore, in endosomes, only receptors undergoing partial proteolysis are able to bind signaling adaptors and build oligomeric signaling complexes (*4*). These signaling complexes induce transcription of the genes coding for type I IFN and proinflammatory cytokines (*5*). For some TLRs, different endosomal populations engage different signaling pathways. Thus, for TLR9, NF-κB-mediated signaling, which triggers the production of proinflammatory cytokines, occurs both in early endosomes described by the SNARE VAMP3 and in lysosomes, while IRF7-mediated signaling occurs only in LAMP1+ lysosomes (*6*). We previously found that the retention of TLR9 and its ligand in VAMP3+ early endosomes is dependent on the presence of IRAP. In the absence of IRAP, TLR9 is constitutively targeted to lysosomes and signaling via TLR9 is enhanced in IRAP-deficient cells and mice (*7*).

Besides TLRs, specific endocytic pathways could regulate the activity of other immune receptors, which are known to be internalized after their activation. We explored the impact of endocytosis on FcRs, a family of immune receptors that play a central role in the inflammatory response. High-affinity receptors, such as Fc∊RI and FcγRI and murine FcγRIV, bind monomeric Igs, while moderate/low-affinity receptors, such as murine FcγRIII and FcγRIV and human FcαRI, FcγRIIA and FcγRIIIA bind only immune complexes (IC). Most FcRs are activating receptors and their signaling is initiated by the phosphorylation of ITAM sequences (immunoreceptor tyrosine-based activatory motifs). In general, the ITAM is provided by the associated γ-chain of FcRs and only the human activating FcγRIIA includes an ITAM-like motif in its own cytosolic tail. After ligand binding, ITAMs are phosphorylated by Src-kinases, which allow the binding of the Syk-kinase to the receptor and the activation of downstream signaling cascades (*8*).

Signaling via activating FcRs is counteracted by the inhibitory receptor FcγRIIB, which contains in its cytosolic tail an immunoreceptor tyrosine-based inhibitory motif (ITIM). When phosphorylated, the ITIM recruits the SHIP and SHP phosphatases that will block the propagation of the signal from an activating receptor (*8*). Although FcR internalization after IC binding is well documented, the intracellular localization of FcR signaling in endosomal platforms has not been investigated yet. In this report, we explored the trafficking of FcγRI, FcγRIIA, FcγRIIB and FcαRI after receptor cross-linking and demonstrated that FcγRI is one of the receptors with the highest ability to build endosomal-signaling platforms, which are IRAP-dependent.

## Results

### Activated FcγRI is retained in intracellular compartments described by IRAP and TGN38

We investigated the trafficking of four different FcRs, one activating IgG receptor (FcγRI), the inhibitory FcγRIIB receptor and two dual function Fc receptors, the FcαRI and the FcγRIIA, which are able to engage via their ITAM an activating signaling triggered by IC, or an inhibitory signaling triggered by monovalent Igs (*9*). The activating IgG receptors were the high affinity IgG receptor FcγRI, paired with the γ-chain, and the low affinity IgG receptor FcγRIIA that does not need the association with the γ-chain for signaling. We used lentiviral infections to express all these human receptors as GFP-fusion proteins in the GM-CSF DC-like murine cell line DC2.4. To evaluate the intracellular trafficking of the receptors, we used the early endosomal antigen 1 (EEA1), which is a marker of early endosomes, the TGN38, a marker of trans-Golgi vesicles (TGN), the lysosome marker LAMP1 and IRAP, the marker of specialized, slow recycling storage endosomes shown to regulate TLR9 signaling (*7*) and more recently antigen T cell receptor (TCR) signaling (*10*). To activate FcγRI, the receptor was crosslinked with anti-FcγRI and its localization was monitored by immunofluorescence. If in basal conditions FcγRI was localized mainly at the plasma membrane, as early as 15 minutes after its crosslinking the receptor was internalized in IRAP^+^ endosomes (Fig.1A) and TGN38^+^ vesicles (Fig.1B). Although we did not directly co-stain IRAP and TGN38, they probably colocalize, as previously demonstrated (*11*, *12*). Interestingly, half of the receptor and the crosslinking antibodies were retained for more than 60 minutes in IRAP-TGN38 endosomes and were transported to LAMP1^+^ lysosomes at later time points (Fig.1C), without significant accumulation in EEA1^+^ endosomes (Fig.1D).

**Figure 1.**
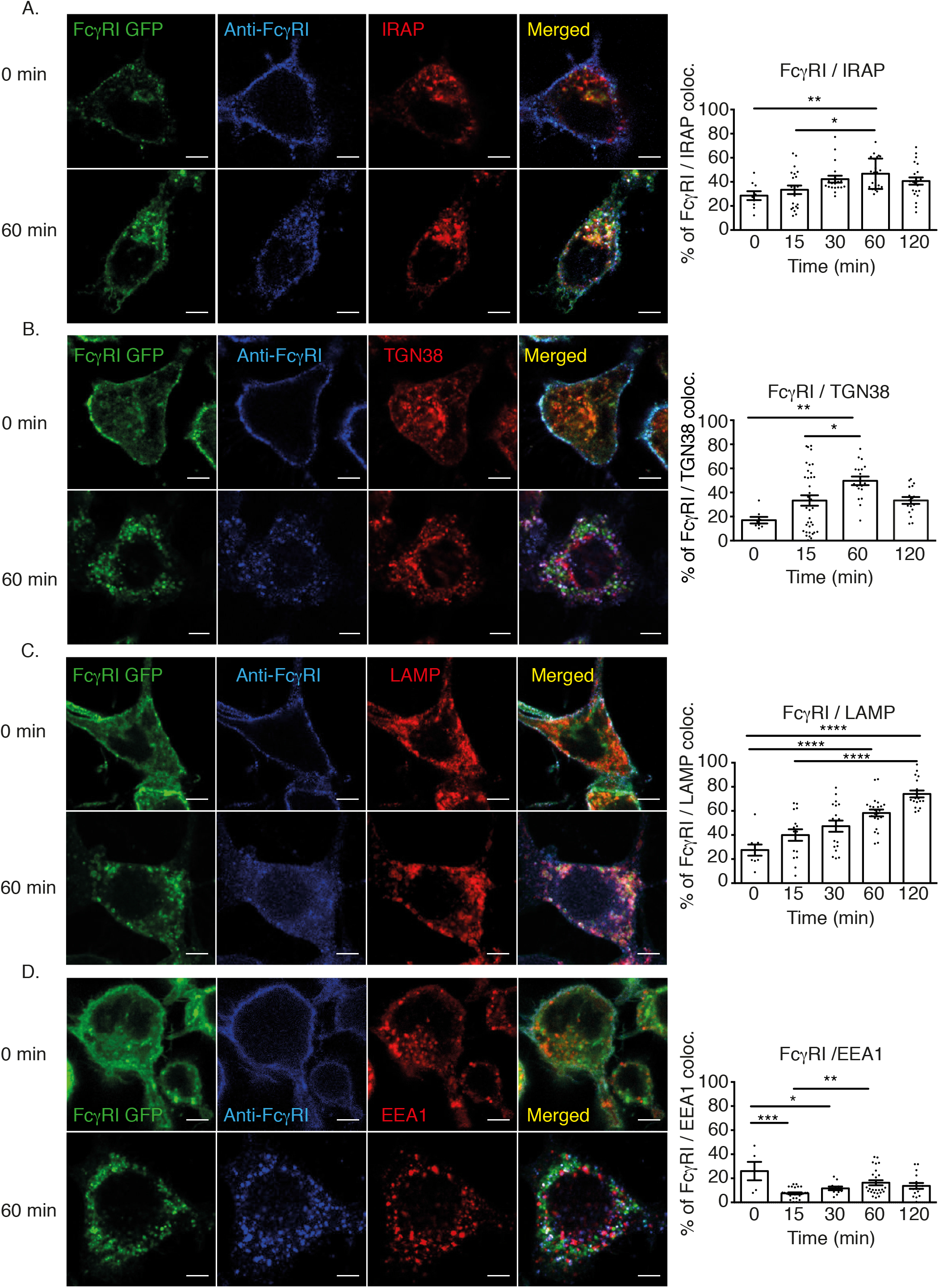
Activated FcγRI is retained in intracellular compartments described by IRAP and TGN38. (A-D) DC2.4 cells expressing FcγRI-GFP (green) were incubated with anti-FcγRI (clone 10.1) and crosslinked with anti-mouse IgG (blue) at 4 °C. After removal of excess antibodies, the cells were shifted for the indicated time points at 37 °C, fixed and stained for IRAP (red) (A), TGN38 (red) (B), LAMP1 (red) (C) and EEA1 (red) (D). The pictures show representative images from three independent experiments and the graphs show colocalization between FcγRI and the endocytic markers. Each dot represents a cell.

### FcγRIIA, FcαR and FcγRIIB are internalized in distinct intracellular compartments

To compare the trafficking of high affinity and low affinity IgG receptors, we performed similar experiments, by crosslinking FcγRIIA with anti-FcγRIIA. Similar to FcγRI, FcγRIIA was internalized in IRAP^+^ endosomes (Fig. 2A), where the receptor-ligand concentration peaked at 30 minutes, and was later targeted to lysosomes (Fig. 2B), probably via EEA1^+^ endosomes (Fig. 2C). We wondered if, similarly to the activating FcγRs, the inhibitory IgG receptor FcγRIIB was targeted to IRAP+ endosomes after internalization. Analysis of trafficking of FcγRIIB crosslinked by anti-FcγRIIB showed that this receptor was found in vesicles containing the autophagy marker LC3, a result that might suggest a new mechanism by which autophagy could dampen the inflammatory response (Fig. S1A-D).

**Figure 2.**
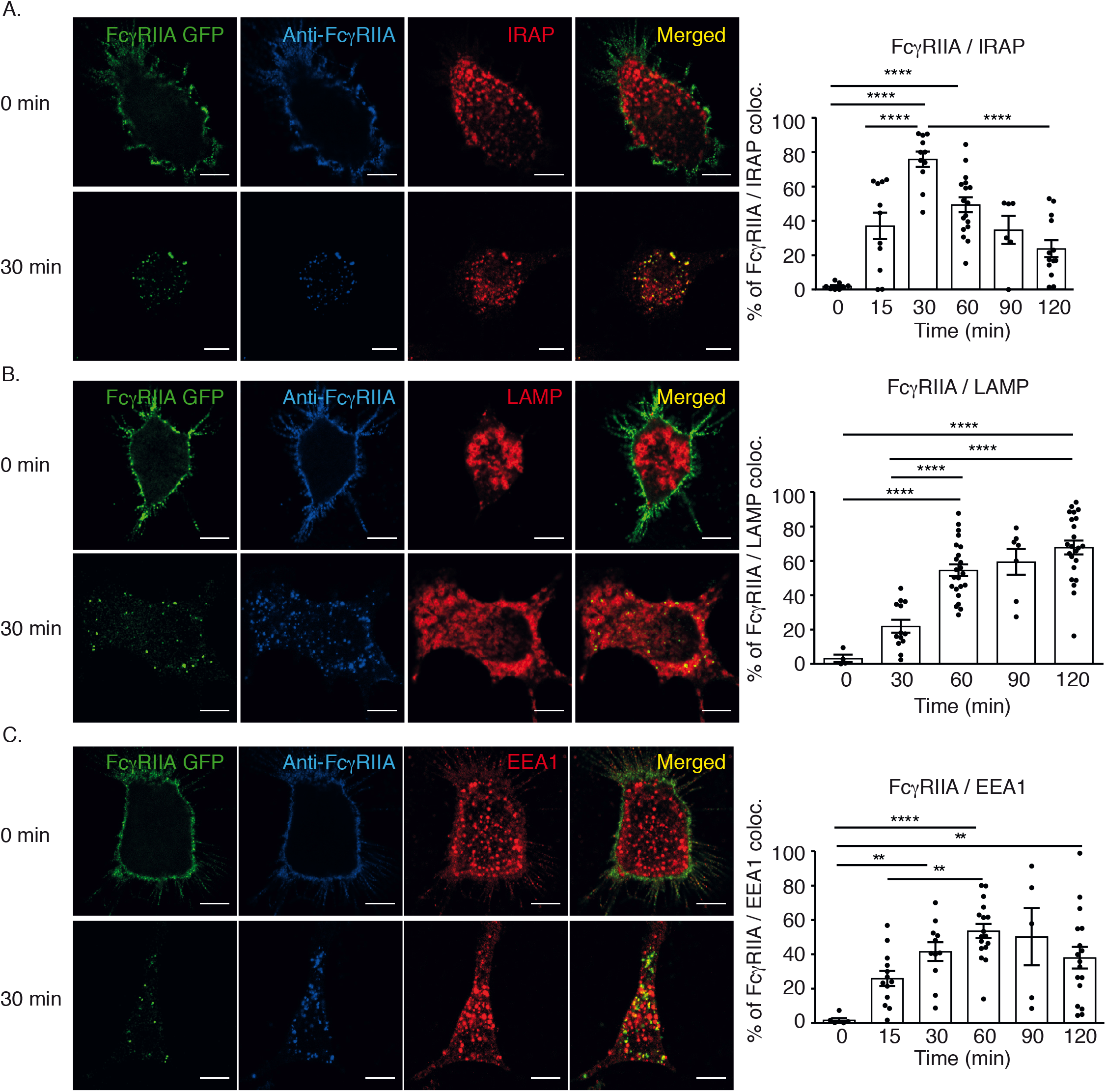
Activated FcγRIIA shows a short passage by IRAP endosomes. (A-C) DC2.4 cells expressing FcγRIIA-GFP (green) were incubated with anti-FcγRIIA (clone IV.3) and crosslinked with anti-mouse IgG (blue) at 4 °C. After removal of excess antibodies, the cells were shifted for the indicated time points at 37 °C, fixed and stained for IRAP (red) (A), LAMP1 (red) (B) and EEA1 (red) (C). The pictures show representative images from three independent experiments and the graphs show colocalization between FcγRIIA and the endocytic markers. Each dot represents a cell. Scale bars = 5 μm.

Since two different activating FcRs were internalized in IRAP^+^ endosomes, we wondered if this feature is specific to IgG receptors, or applies to all activating FcRs. To answer this question, we investigated the trafficking of the FcαRI receptor after crosslinking with the specific antibody A77 (*13*). In contrast with the behavior of both IgG receptors, FcαRI accumulated much less in IRAP vesicles and was poorly internalized in lysosomes (Fig. S2A-C). In conclusion, among the four receptors investigated, the activating FcγRs and the corresponding crosslinking antibodies were internalized at a higher extent in IRAP^+^ vesicles, where FcγRI was retained for a longer time than FcγRIIA.

### Activated FcγRI recruits active Syk in IRAP endosomes

Intrigued by this long retention of FcγRI in IRAP endosomes, we wondered if the receptor is able to signal from this location and we investigated the colocalization between the receptor and the signaling kinase Syk, in basal conditions and after cell stimulation by receptor cross-linking with human IgG and anti-human IgG. In basal conditions, 20 to 40% of total Syk colocalized with FcγRI, mainly at the plasma membrane, but also in small endosomes labeled by IRAP (Fig. 3A).

**Figure 3.**
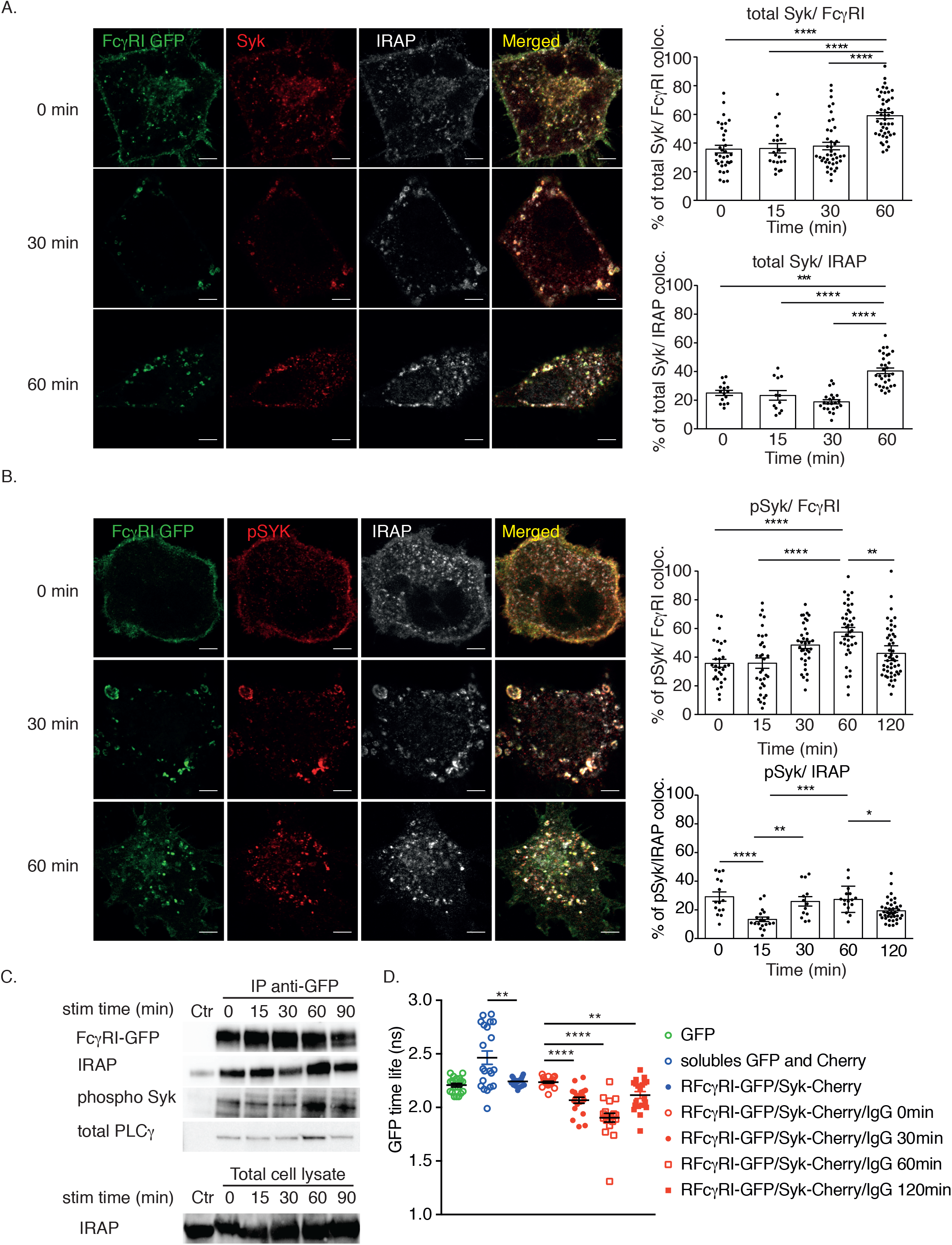
Activated FcγRI recruits active Syk in IRAP endosomes. (A-B) DC2.4 cells expressing FcγRI-GFP (green) were incubated with human IgG and anti-human IgG at 4 °C. After removal of excess antibodies, cells were shifted for the indicated time points at 37 °C, fixed and stained for Syk (red) and IRAP (gray) (A), or Syk phosphorylated on the Tyrosine 352 (pSyk) (red) and IRAP (gray) (B). The pictures show representative images from three independent experiments and the graphs show colocalization between FcγRI or Syk and the endocytic marker, IRAP. Each dot represents a cell. Scale bars = 5 μm. (C) FcγRI-GFP was immunoprecipitated with anti-GFP beads from DC2.4 cells activated as in (A-B) and the immunoprecipitated material was analyzed by immunoblot using the indicated antibodies. The blots are representative of 2 independent experiments. (D) The interaction of FcγRI with Syk was measured by FLIM experiments and the graph show the GFP lifetime for cells expressing only GFP, both GFP and mCherry as soluble proteins and the cells expressing both FcγRI-GFP and Syk-mCherry in steady state and after the receptor crosslinking for indicated time points. Each dot represents a cell.

The basal Syk colocalization with FcγRI was expected, since FcγRI is able to bind with high affinity monomeric bovine IgG from the cell culture medium. When the cells were cultured with medium containing IgG-depleted bovine serum, the receptor recruited Syk only after its stimulation by cross-linking (Fig. S3A-B). However, we used culture medium containing IgG, since the presence of monovalent IgG is the physiological situation and it is known that after receptor crosslinking, FcγRI preferentially interacts with ICs, in spite of monomeric IgG pre-engagement (*14*). Indeed, after activation by receptor crosslinking, the FcγRI was massively internalized in IRAP^+^ vesicles, which fused to create enlarged endosomes to which Syk was recruited (Fig. 3A). Recruitment of Syk to the double phosphorylated ITAM of FcγRs requires an initial phosphorylation of ITAMs by Src-kinases. Once bound to phosphorylated ITAMs, Syk changes its conformation to an open and active form. Active Syk is able to autophosphorylate itself on multiple tyrosine residues that will allow direct recruitment of several proteins, such as class I PI3K, PLCγ, VAV and SLP76, associated with FcγR downstream signaling. In addition to its autophosphorylation, active Syk is also able to phosphorylate the ITAM tyrosine of the receptor, providing thus a positive feedback loop for FcγR signaling (*8*). To see if endosomal Syk is phosphorylated, we stained the cells with an antibody recognizing the phosphorylated Tyrosine 352 of active Syk. Similar to total Syk, active Syk was in basal conditions at the plasma membrane and in small endosomes, but after receptor crosslinking more than half of the active Syk and FcγRI, were found in IRAP endosomes, where they peaked at 60 minutes after activation (Fig. 3B). These results were confirmed by co-immunoprecipitation experiments showing that FcγRI pulled down the highest amounts of IRAP, pSyk and PLCγ at 60 minutes (Fig. 3C).

An alternative method to investigate protein interaction is the FRET-FLIM (Fluorescence Resonance Energy Transfer – Fluorescence Lifetime IMaging). To investigate FcγRI interaction with Syk by FRET, we expressed FcγRI-GFP and Syk-mCherry in DC2.4 cells (Fig. S3C) and measured the lifetime of GFP in steady state and after receptor crosslinking. The minimal GFP lifetime, which is equivalent to the highest interaction between FcγRI-GFP and Syk-mCherry, was observed at 60 minutes after the receptor crosslinking (Fig. 3D and Fig. S3D). These results suggest that IRAP storage endosomes could act as endosomal signaling platforms for the high affinity IgG receptor, FcγRI. This receptor is also activated by monomeric IgG (*15*), activation confirmed by our experiments in which the cell culture with bovine serum IgG caused a basal recruitment of active Syk to the receptor, mainly at the plasma membrane (Fig. 3B). When a polyvalent immune complex was used to trigger FcγRI, active Syk remained associated with the receptor in IRAP endosomes as long as 1 hour after receptor activation. The endocytosis of FcγRI activated by polyvalent ICs might therefore be used to discriminate between a monovalent and a polyvalent ligand.

### IRAP deletion affects the endosomal recruitment of pSyk to FcγRI

Considering the previously demonstrated role of IRAP in storage endosome trafficking (*16*) and TLR9 (*7*) and TCR activation (*10*), we wanted to know if, similar to TLR9 and TCR, FcγRI trafficking or signaling was affected in the absence of IRAP. Since the DC2.4 cell line did not survive when we depleted IRAP, we used bone marrow derived dendritic cells (BM-DCs) obtained from wild type and IRAP-deficient mice (*17*) to investigate the trafficking of both chains of endogenous FcγRI, the chain encoded by *Fcgr1* gene and the common FcR-γ-chain associated with FcRs, encoded by the murine gene *Fcer1g* (*18*). While in wt BM-DCs CD64 colocalized in endosomes with the FcR-γ-chain as long as 2 hours after the receptor crosslinking with the anti-murine Fcgr1 specific antibody, in the absence of IRAP this colocalization was lost at 2 hours and both Fcgr1 and the FcR-γ-chain showed a weak to undetectable staining (Fig. 4A). This suggests that in the absence of IRAP, both chains of activated FcγRI are targeted to degradation. However, at early time points, when the FcR-γ-chain was still detectable in IRAP-deficient BM-DCs, we observed that the colocalization between the FcR-γ-chain and active Syk was reduced in the absence of IRAP (Fig. 4B), indicating that IRAP is required for the stabilization of endosomal signaling platforms of endogenous murine FcγRI.

**Figure 4.**
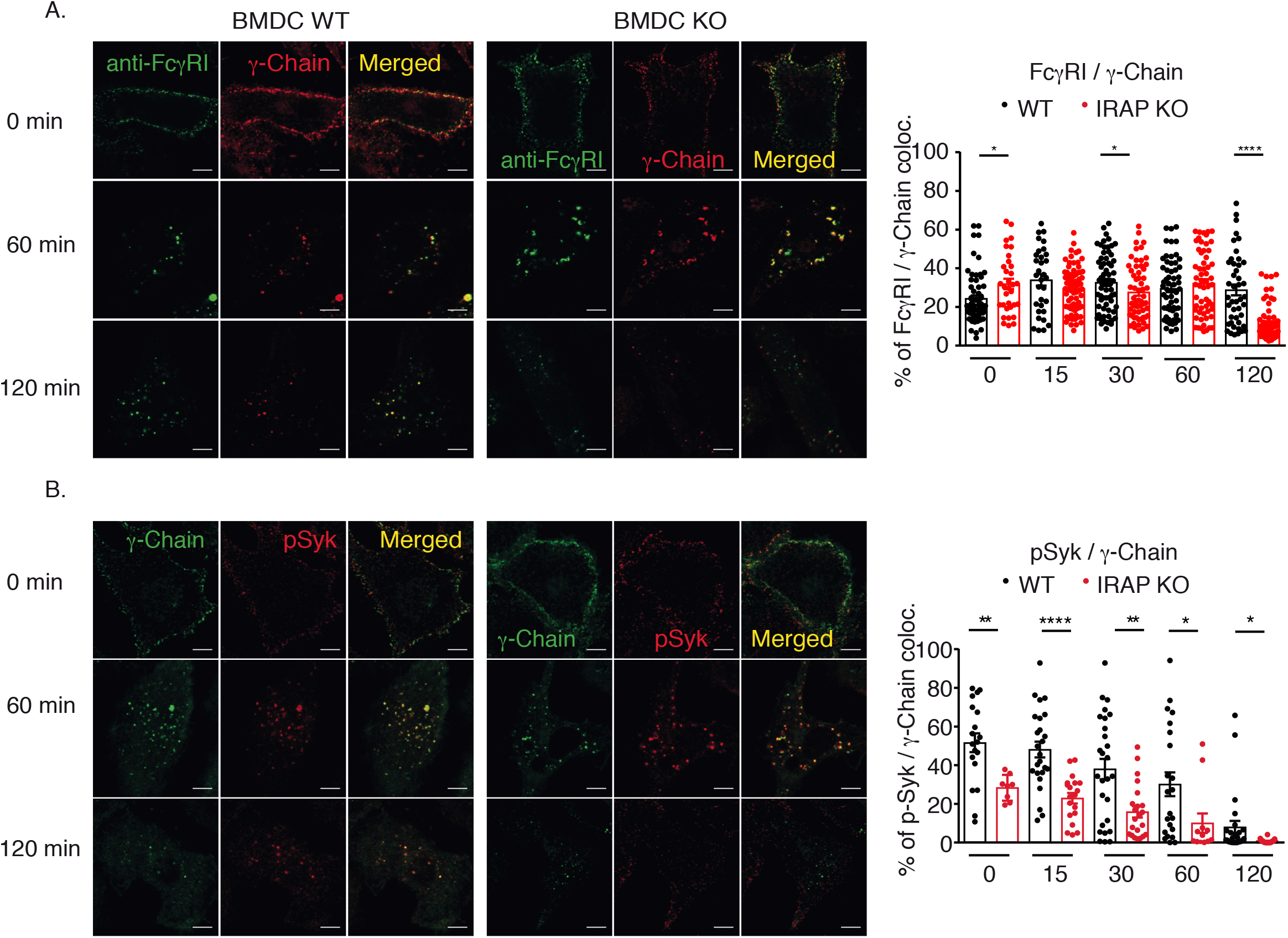
IRAP deletion affects the endosomal recruitment of active Syk to FcγRI. (A-C) Wild-type (WT) or IRAP-deficient (KO) BM-DCs were incubated with anti-mouse FcγRI (clone AT152-9), followed by crosslinking with non-labeled anti-rat IgG at 4 °C. After removal of excess antibodies, the cells were shifted for the indicated time points at 37 °C, fixed and stained for endogenous FcγRI (green) and γ chain (red) in (A) and for γ chain (green) and Syk phosphorylated on Tyrosine 352 (pSyk) (red) in (B). The pictures show representative images from three independent experiments and the graphs show colocalization between FcγRI and the γ chain (A) and colocalization between the γ chain and phosphorylated, active Syk (B). Each dot represents a cell. Scale bars = 5 μm.

The major small GTPase recruited on IRAP^+^ endosomes in BM-DCs is Rab14 (*19*) and we have previously shown that Rab14 interacts with the kinesin KIF16B. The Rab14-KIF16B complex is involved in the anterograde transport of IRAP endosomes along microtubules and in their fusion with early endosomes and newly formed phagosomes (*16*). Since Rab14 depletion destabilized IRAP endosomes, we wondered if FcγRI endosomal signaling would also be affected in the absence of Rab14. To investigate this, we depleted Rab14 in DC2.4 cells using specific lentiviral shRNA (Fig. S4A) and investigated FcγRI colocalization with IRAP and active Syk. In the absence of Rab14, FcγRI colocalization with IRAP was lower than in control cells, due to the decreased formation of IRAP^+^ endosomes (Figure S4B), in agreement with our previously published results (*16*). In parallel, the amount of pSyk recruited to FcγRI was significantly decreased in Rab14 depleted cells (Fig. S4C). These results show that Rab14 is also involved in the sustained endosomal signaling of FcγRI. Considering that Rab14 transports IRAP^+^ endosomes to the cell periphery and decreases their fusion with lysosomes by interacting with KIF16B in GMCSF derived BM-DCs (*16*), it is expected that in the absence of Rab14, activating FcγRs, that are cargos of IRAP endosomes, will be transported to lysosomes and degraded. Similar mechanisms were demonstrated for EGFR in Hela cells, where KIF16B activity prevents EGFR targeting to lysosomes, thus facilitating receptor recycling and signaling (*20*). Our results indicate that similar to growth factor receptors, activating FcγRs, and especially the high affinity receptor FcγRI, use not only the plasma membrane for signaling, but also specific endosomal compartments, which are described by IRAP and Rab14.

### IRAP-deficient macrophages show reduced ADCC

To investigate the impact of IRAP deletion on activating FcγR functions (*21*), we performed an antibody dependent cellular cytotoxicity (ADCC) assay, which estimates the efficiency of FcγR signaling (*22*). We used wt and IRAP-deficient macrophages (MF) as effector cells, the HUCCT1 cholangiocarcinoma cell line expressing EGFR as target cells and an anti-EGFR antibody to trigger the IgG2a-mediated cell cytotoxicity, a phenomenon in which FcγRI has a predominant role (*23*). After 2 days incubation of MFs with HUCCT1 cells, in the presence of increasing anti-EGFR antibody concentrations, we observed that the killing of tumor cells by IRAP-deficient MFs was less efficient than the killing by wt MF (Fig. 5). This impaired killing of HUCCT1 targets by IRAP-deficient MF was not due to a reduction in the cell surface expression of the FcγRI (Fig. S5), suggesting that endosomal signaling is required for FcγR ability to trigger ADCC.

**Figure 5.**
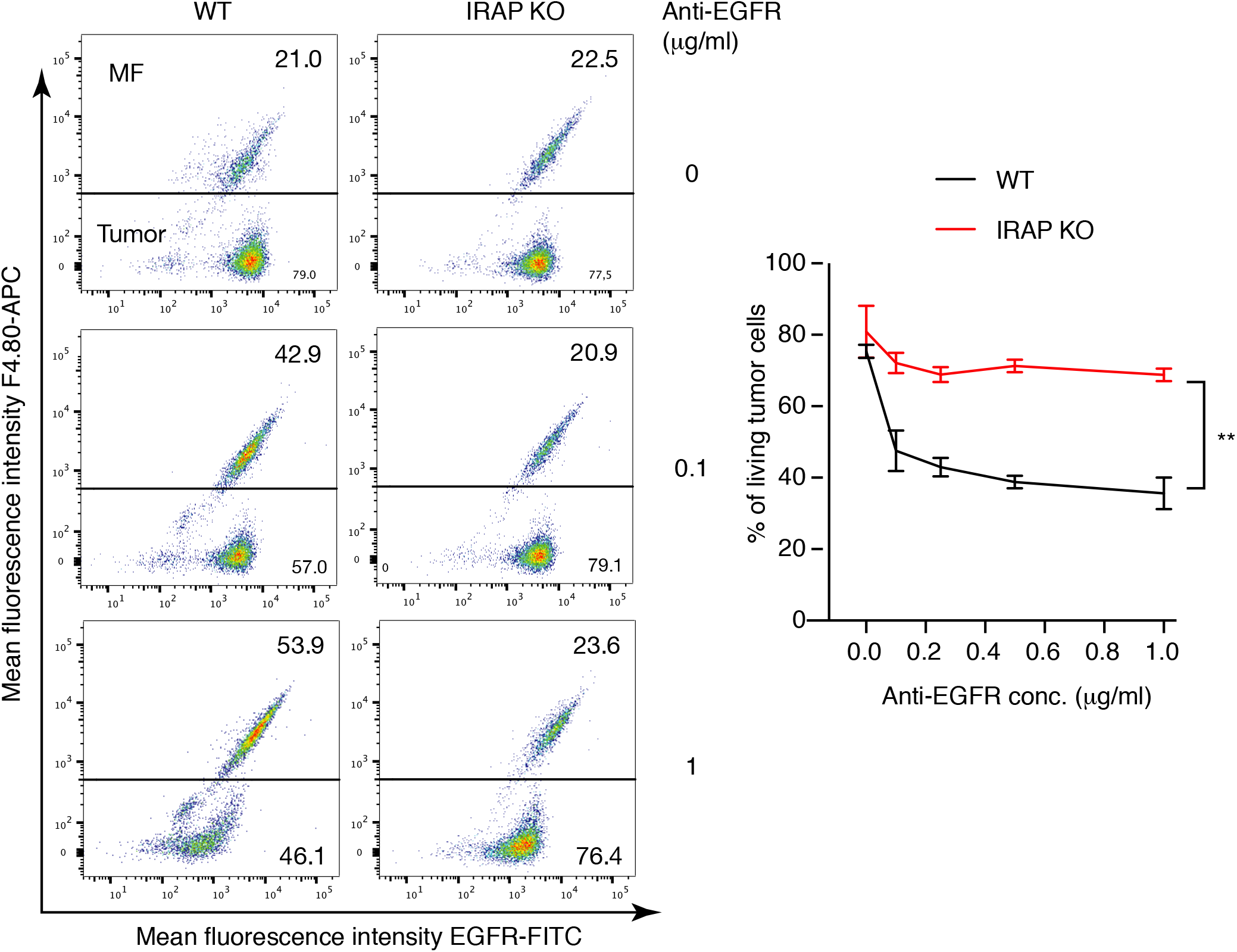
IRAP-deficient macrophages show reduced ADCC. Wild-type (WT) or IRAP-deficient (KO) peritoneal MF were incubated with HUCCT1 tumor cells expressing EGFR in the presence of indicated concentrations of mouse IgG2a anti-EGFR antibody. After 3 days the cells were stained with anti-F4/80 to identify MFs and with mouse IgG1 anti-EGFR, followed by FITC conjugated goat F(ab’)2 anti-mouse IgG1 to identify the tumor cells. The pictures show representative images from three independent experiments and the graph shows the percentage of living HUCCT1 cells defined as EGFR positive and F4/80

## Discussion

Our study showed that activating FcγRs, similar to non-immune cell surface receptors such as RTKs (*2*) are able to use endosomal-signaling platforms. Both in RTK and GPCR signaling, the targeting of the ligand-receptor complexes towards degradation or towards signaling endosomes depends on the nature of the ligand. Thus, at low ligand concentrations, EGFR is internalized via clathrin dependent endocytosis (CDE), while at high ligand concentrations EGFR is internalized through both CDE and clathrin independent endocytosis (CIE). EGFR endocytosis via CDE leads to increased signaling, while the CIE pathway directs the receptor to lysosomes, leading to receptor degradation and limiting EGFR signaling. The exact sub-pathway of CIE that is used by EGFR is not yet identified (*24*). Interestingly, the endocytic pathways that transport the receptors in signaling endosomes are specific for each type of receptor. Thus, LRP6 (low-density lipoprotein receptor-related protein 6), a receptor connected to Wnt signaling, use the two pathways of endocytosis in a way that seems the reverse of what was observed for EGFR. CIE of LRP6 leads to sustained signaling, while LRP6 degradation is a consequence of the receptor internalization by CDE (*25*).

Similar to the previously mentioned receptors, such mechanisms might apply to FcγRs. Pioneer studies of FcR signaling and trafficking have shown that FcγR activation by polyvalent ligands induces FcR internalization and their final targeting to lysosomes (*13*, *26*), while FcγR ligation by monovalent IgGs induces the recycling of the receptor (*27*, *28*). These initial studies did not address the pathway by which the FcγR is internalized in the different activation scenarios or the precise intracellular compartments that are involved.

Even if our study did not identify the endocytic pathway that leads to activating FcRs internalization in IRAP^+^ signaling endosomes, we demonstrated that the FcγRs activation by polyvalent ligands triggers an endocytosis mechanism required for a sustained FcγR signaling from endosomal vesicles described by IRAP and Rab14. All our experiments were performed using receptor cross-linking, which is mimicking polyvalent IC ligands, a situation in which ITAM-coupled FcRs function as pro-inflammatory receptors. However, ITAM-coupled FcRs bind also monovalent ligands, in which case, they act as anti-inflammatory receptors (*29*). FcγR binding to monovalent IgGs induces a partial phosphorylation of ITAM motif, followed by the recruitment of SHIP phosphatases and an inhibitory signaling, usually called ITAMi, for inhibitory ITAM (*30*). This ITAMi signaling is able to control the amplitude of activation of several other immune receptors, including TLRs and FcRs (*29*, *30*), being thus, a key factor of immune homeostasis, in addition to the inhibitory signaling provided by ITIM-coupled immune receptors.

Whether ITIM-coupled receptors and FcRs associated ITAMi use the same signaling platforms is not yet known. The ITAMi signaling of activating FcR was shown to induce accumulation of both activated and inhibitory receptors in an “intracellular cluster” (*31*), but the identity of this “intracellular cluster” has not yet been identified. We show here that the internalization of the ITIM-coupled inhibitory receptor FcγRIIB occurs in LC3^+^ vesicles, raising the question of whether all inhibitory signals of FcRs are delivered from LC3^+^ vesicles, which are probably autophagosomes. Autophagy was reported as an anti-inflammatory pathway via at least two mechanisms: the elimination of a plethora of damaged organelles released stimuli that activate the inflammasome and by the decrease of pathogen load that will diminish the innate immune receptors activation (*32*). Building of ITIM and ITAMi signaling platforms on autophagosomes, which should be investigated, might add another mechanism explaining previously reported anti-inflammatory functions of autophagy.

In conclusion, our study that analyzed intracellular FcγR trafficking and signaling only after polyvalent FcγR activation, demonstrated that activating FcγR are able to signal from endosomal platforms and this endosomal signaling is important, at least in MFs, for ADCC, a major mechanism of FcRs signaling that is at the heart of anti-tumoral monoclonal antibody therapies. In addition to ADCC and immunotherapies, this previously unappreciated feature of FcγR signaling might be important in the context of the physiological maintenance of immune homeostasis and in FcγR-mediated diseases physiopathology (*33*). Finally, our results on endosomal signaling of FcRs raise the question whether other ITAM coupled receptors might use endosomes as signaling platforms, like it has been shown for the B cell receptor (*34*) and T cell receptor (*10*, *35*).

## Materials and Methods

### Mice

Previously described IRAP deficient mice (*17*) were bred in our animal facility in pathogen free conditions. All animal care and experimental procedures were performed in accordance with the guidelines and regulations of the French Veterinary Department and approved by the “Comité d’Ethique pour l’Experimentation Animale” of our institute.

### Cell culture

BM-DCs were produced *in vitro* by culturing bone marrow precursor cells from large bones for 7 days in complete medium (IMDM complemented with 10% FCS, 2 mM glutamine, 100 U/mL penicillin, 100 μg/mL streptomycin, 50 μM β-mercaptoethanol) supplemented with 20 μg/mL GM-CSF). DC2.4 were cultured in complete medium (IMDM complemented with 10% FCS, 2 mM glutamine, 100 U/mL penicillin, 100 μg/mL streptomycin, 50 μM β-mercaptoethanol).

### Plasmid constructs and cell transfection

The cDNA for different FcRs was cloned in fusion with EGFP in the pLVX-CMV-IRES-Puro plasmid (Clontech). The cDNA coding for FcγRI was amplified from the NM_000566 cDNA (Origene) with the following primers: forward 5’-TACTCGAGATGTGGTTCTTGACAACTCT -3’ (contains the XhoI restriction site) and reverse 5’-ATGGGCCCCCGTGGCCCCCTGGGGCTC-3’ (contains the ApaI restriction site), cloned into the pCRBlunt shuttle vector (Invitrogen), sequenced and transferred into pEGFP-N1 using the XhoI and ApaI restriction enzymes. Finally, the FcγRI-GFP fusion cDNA was removed from the pEGFP-N1 vector with NheI (blunted with Klenow) and NotI and inserted into pLVX-CMV-IRES-Puro between EcoRI (blunted with Klenow) and NotI.

The cDNA coding for FcγRIIA-GFP in pLCPX vector was a kind gift from Peter Cresswell (*36*). FcγRIIA-GFP was removed from pLPCX with BglII (blunted with Klenow) and NotI and cloned into pLVX-EF1a-IRES-Puro between EcoRI (blunted with Klenow) and NotI.

The FcαRI-GFP in pSelect and the cDNA coding for FcγRIIB were provided by Erwan Boedec (INSERM U1149, Paris). The RFcαI-GFP fusion cDNA was amplified with the following primers: forward 5’-TAgaattcATGGACCCCAAACAGACCac-3’ (contains the EcoRI restriction site) and reverse 5-TAgcggccgcTTACTTGTACAGCTCATCCATTCC-3’ (contains the NotI restriction site). The PCR product was cloned into pCRBlunt (Invitrogen). After sequencing, the cDNA coding for FcαRI-GFP fusion was transferred into pLVX-CMV-IRES-Puro between EcoRI and NotI.

The FcγRIIB cDNA was amplified with the following primers: forward 5’-TAACCGGTATGGGCATACTGAGCTTTCT-3’ (contains the AgeI restriction site) and reverse 5’-TAGGATCCAATCCTGTTCTGGTCGTCG-3’ (contains the BamHI restriction site) and the PCR product was cloned into pCRBlunt (Invitrogen) and sequenced. From pCRBlunt, the FcγRIIB cDNA was transferred into the pSelect-GFP-C vector using AgeI and BamHI. Finally, the FcγRIIB-GFP fusion cDNA was amplified with the following primers: forward 5‘-TAGAATTCATGGGCATACTGAGCTTTCT-3’ (contains the EcoRI restriction site) and reverse 5’-TAgcggccgcTTACTTGTACAGCTCATCCA-3’ (contains the NotI restriction site) and cloned into pLVX-CMV-IRES-Puro between EcoRI and NotI.

### Antibodies

Primary antibodies used were: anti-human-RFcγI mouse monoclonal antibody, clone 10.1 (BioRad), anti-IRAP rabbit monoclonal antibody, clone D7C5 (Cell Signaling), anti-TGN38 rabbit polyclonal antibody (Thermo Fisher Scientific), anti-LAMP1 rat monoclonal antibody, clone 1D4B (Thermo Fisher Scientific), anti-EEA1 goat polyclonal antibody (Santa Cruz, N-19), anti-LC3B rabbit polyclonal antibody (Cell Signaling), anti-Syk rabbit monoclonal antibody (Cell Signaling), anti-phospho-Syk (Tyr352), clone 65E4 (Cell Signaling), anti-γ chain mouse monoclonal antibody (MBL) and anti-Rab14 rabbit polyclonal serum (Sigma).

Secondary antibodies used were: AlexaFluor-647 conjugated donkey anti-rabbit (Invitrogen), AlexaFluor-405 conjugated donkey anti-rabbit, donkey anti-rat or donkey anti-goat (Abcam) and AlexaFluor-594- or AlexaFluor-633-conjugated donkey anti-mouse (Invitrogen).

### Lentivirus Production and Transduction

Viruses were produced using standard protocols (*37*), using the previously published shRNA in pLKO.1 plasmids (*16*) and the packaging plasmids pCMVDelta8.2 and pMD2G. Lentiviral particles were produced in HEK293-FT cells (Invitrogen). Then, DC2.4 cells were seeded in 96-well flat-bottom plates at 2 × 10^4^ cells per well and transduced in the presence of 10 μg/ml polybrene. Following 90 min centrifugation at 37 °C and 950×g, the lentiviral mix was replaced with complete IMDM medium. The day after, puromycin was added to the cells at 5 μg/ml and selected cells were used 4-5 days post transduction.

### Receptor aggregation for the microscopy assays

DC2.4 cells or BM-DCs were seeded on fibronectin-coated slides for 16 h. To aggregate specific Fcγ receptors, cells were incubated with specific monoclonal antibodies for 15 min on ice. To aggregate FcγRI, the cells were incubated with 5 μg/ml of mouse anti-human CD64, clone 10.1 (Biolegend). To aggregate FcγRI mouse on BM-DCs, cells were incubated with 15 μg/ml of rat anti-mouse CD64, clone AT152-9 (BioRad). To aggregate FcγRIIA, cells were incubated with 10 μg/ml of mouse anti-human FcγRIIA, clone IV.3 (Stemcell). To aggregate FcγRIIB, cells were incubated with 10 μg/ml of mouse anti-human FcγRIIB, clone AT10 (Santa Cruz). To aggregate FcαRI, cells were incubated with 10 μg/ml of F(ab)’2 fragment of the monoclonal mouse antibody A77. After removal of excess antibody by two washes in PBS, the receptors were crosslinked by the addition of anti-mouse or anti-rat IgG Alexa Fluor (Sigma) during 15 min on ice.

For experiments using IgGs to aggregate the receptors, cells were incubated for 15 min on ice with 5 μg/ml human polyclonal IgG (ChromPure, Jackson ImmunoResearch). Unbound IgGs were removed by two washes in cold PBS. The cells were then incubated with 10 μg/ml of donkey anti-human IgG (Jackson ImmunoResearch) on ice for 15 min, to aggregate the receptors. Cells were then shifted to 37 °C for the time points indicated in the assays.

### Immunofluorescence microscopy

For all experiments cells were fixed with 2% PFA for 15 min at 37 °C, permeabilized with 0.2% saponin in PBS containing 0.2% BSA and stained in the same buffer. Images were acquired on a Leica SP8 confocal microscope. Image treatment and analysis were performed with ImageJ software.

### Receptor aggregation for the biochemical assays (WB, Co-IP)

Cells were incubated for 15 min on ice with 5 μg/ml human polyclonal IgG (ChromPure, Jackson ImmunoResearch) and the excess of unbound IgGs was removed by two washes in PBS. The receptors were then aggregated by addition of 10 μg/ml of donkey anti-human IgG (Jackson ImmunoResearch) for 15 min on ice. The excess of unbound antibodies was removed by two washes in PBS and the cells were shifted to 37 °C for the different time points of the assays.

### Co-immunoprecipitation and Immunoblots

Cell pellets were lysed in 50mM Tris, 150mM NaCl, 1% CHAPS supplemented with Complete protease inhibitor mix (Roche) and phosphatase inhibitor cocktails 2 and 3 (Sigma-Aldrich). Lysates were cleared by 10 min centrifugation at 15000xg and incubated with GFP-trap beads (Chromotek). The material bound on the beads was eluted in Laemmli buffer and the proteins were separated by SDS-PAGE using Criterion 4-15% acrylamide gels (BioRad) in Tris-Glycine-SDS buffer. The proteins were transferred on PVDF membranes (BioRad) using a Trans-Blot® Turbo™ Transfer System from BioRad. Membranes were blocked overnight in 4% non-fat milk and incubated 1h or overnight with each antibody, washed extensively and incubated for 5 min with Clarity™ Western ECL Substrate (BioRad). The chemiluminescence signal was acquired using a ChemiDoc™ Imaging System and the quantification was realized with the ImageLab software (BioRad).

### FLIM (Fluorescence Lifetime Imaging Microscopy)

The pMSCV-mCherry-Syk plasmid (Addgene no. 50045) (*38*) was used to express via a retrovirus mCherry-Syk in DC2.4 expressing the FcγRI-GFP. The cells were selected in puromycine and cloned by limiting dilutions.

The FcγRI-GFP receptors were aggregated using the same protocol as for the co-immunoprecipitation experiments. After the removal of the unbound antibodies, the cells were shifted to 37 °C for the different time points of the assays, fixed and analyzed for FRET measurements. Images were acquired on a TCS SMD FLIM Leica SP8 confocal microscope and SMD PicoQuant. Image treatment and analysis were performed with Software Symphotime.

### ADCC

Wt and IRAP-deficient mice were injected intra-peritoneal with 1 ml of 3.8% thiolycollate medium and the peritoneal cells were harvested 72 h later using cold PBS. The MF number was estimated by flow cytometry using the F4/80 antibody and plated on 24 wells plates at 0.2 x106 F7/80+ cells per well. Sixteen hours later 0.1 x106 HUCCT1 cells were added to each well in the presence of increasing concentrations of anti-EGFR antibody (mouse IgG2a, clone 528, Invitrogen). Three days later, the cells were stained with F8/40-APC and anti-EGFR antibody (mouse IgG1, clone 225, ThermoFisher Scientific), followed by goat F(ab’)2 anti-mouse IgG1-FITC (Southern Biotech). The percentage of living HUCCT1 cells was estimated by flow cytometry, as EGFR^+^, F4/80^−^ cells.

### Statistical Analysis

Values are expressed as mean ± SEM. Statistical significance between two groups was analyzed using the unpaired Student’s t test, p values are indicated as: *p < 0.05; **p < 0.01; ***p < 0.001; ****p < 0.0001. GraphPad Prism version 6.0 was used to perform the statistical analysis.

## Acknowledgments

We thank the Labex Inflamex and the Agence Nationale pour la Recherche (ANR grants IDEA, ECLIPSE and Help2Kill) for financial support. We also thank Meriem Garfa-Traore of the Imaging Core Facility of SFR Necker for helping with FLIM-FRET analysis and Gregory Gauthier for peritoneal macrophage isolation.

## Author Contributions

Conceptualization: L.S and S.B; Methodology: Most experiments were performed by S.B, with help from M.N and M.B, I.E and R.M helped with advice on experiments and contributed to critical reading of the manuscript, M.C. D. L. performed a part of molecular cloning; Writing - original draft: L.S & S.B., revised version- I.E and L.S; Funding acquisition: L.S.

## Supplementary Information

**Fig. S1. After crosslinking, FcγRIIB is internalized in endosomes containing the autophagy marker, LC3**

(A-D) DC2.4 cells expressing FcγRIIB-GFP (green) were incubated with mouse anti-human Fc◻RIIB (clone AT10) and crosslinked with anti-mouse IgG (blue) at 4 °C. After removal of excess antibodies, the cells were shifted for the indicated time points at 37 °C, fixed and stained for EEA1 (red) (A), IRAP (red) (B), LAMP1 (red) (C) and LC3 (red) (D). The pictures show representative images from three independent experiments and the graphs show colocalization between FcγRIIB and the endocytic markers. Each dot represents a cell. Scale bars = 5 μm.

**Fig. S2. After crosslinking, FcαRI is poorly internalized and targeted to lysosomes**

(A-C) DC2.4 cells expressing FcαRI-GFP (green) were incubated with mouse anti-human FcαRI (clone A77) and crosslinked with anti-mouse IgG (blue) at 4 °C. After removal of excess antibodies, the cells were shifted for the indicated time points at 37 °C, fixed and stained for EEA1 (red) (A), IRAP (red) (B), and LAMP1 (red) (C). The pictures show representative images from three independent experiments and the graphs show colocalization between FcαRI and the endocytic markers. Each dot represents a cell. Scale bars = 5 μm.

**Fig. S3. In the absence of bovine IgG in the cell culture, FcγRI does not recruit Syk in basal conditions**

(A) DC2.4 cells expressing FcγRI-GFP (green) were incubated with human IgG and anti-human IgG at 4 °C. After removal of excess antibodies, the cells were shifted for the indicated time points at 37 °C, fixed and stained for pSyk (red) and IRAP (blue). The pictures show representative images from 2 independent experiments. Scale bars = 5 μm. (B) Quantification of colocalization between the indicated proteins, for the microscopy images obtained in (A). Each dot represents a cell. (C) Representative images of DC2.4 cell lines expressing FcγRI-GFP and Syk-mCherry fusion proteins at 0 and 60 min after the receptor crosslinking by human IgG anti anti-human IgGs. (D) Representative images of FRET-FLIM signal showing GFP average lifetime in DC2.4 cells activated by IgG/anti-IgG ICs.

**Fig. S4. Rab14 is required for the stability of FcγRI-Syk association in IRAP endosomes**

(A) DC2.4 cells expressing FcγRI-GFP were transduced with control or Rab14 shRNA lentiviral particles and Rab14 depletion was analyzed by immunoblotting. (B-C) The cells transduced with control or Rab14 shRNA were incubated with mouse anti-human FcγRI (clone 10.1) and crosslinked with unlabeled anti-mouse IgG. After removal of excess antibodies, the cells were shifted for indicated time points at 37 °C, fixed and stained for IRAP (gray) and pSyk (red) (B), and for EEA1 (gray) and pSyk (red) (C). The pictures show representative images of Rab14-depleted cells. The graphs show colocalization between FcγRI and IRAP (upper graph) and FcγRI and pSyk (lower graph), in both control and Rab14 depleted cells. Each dot represents a cell. Scale bars = 5 μm.

**Fig. S5. IRAP deletion does not affect the cell surface expression of FcγRI**

Macrophages were isolated from the peritoneal cavity of wt and IRAP-deficient mice, stained with anti-FcγRI-PE (clone X54-5/7.1) and analyzed by flow cytometry. The histograms show the anti-FcγRI antibody fluorescence intensity.

