## Supplementary figures and images for "High affinity FcγR activating function depends on IRAP^+^ endosomal-signaling platforms"

### Supplementary Fig1

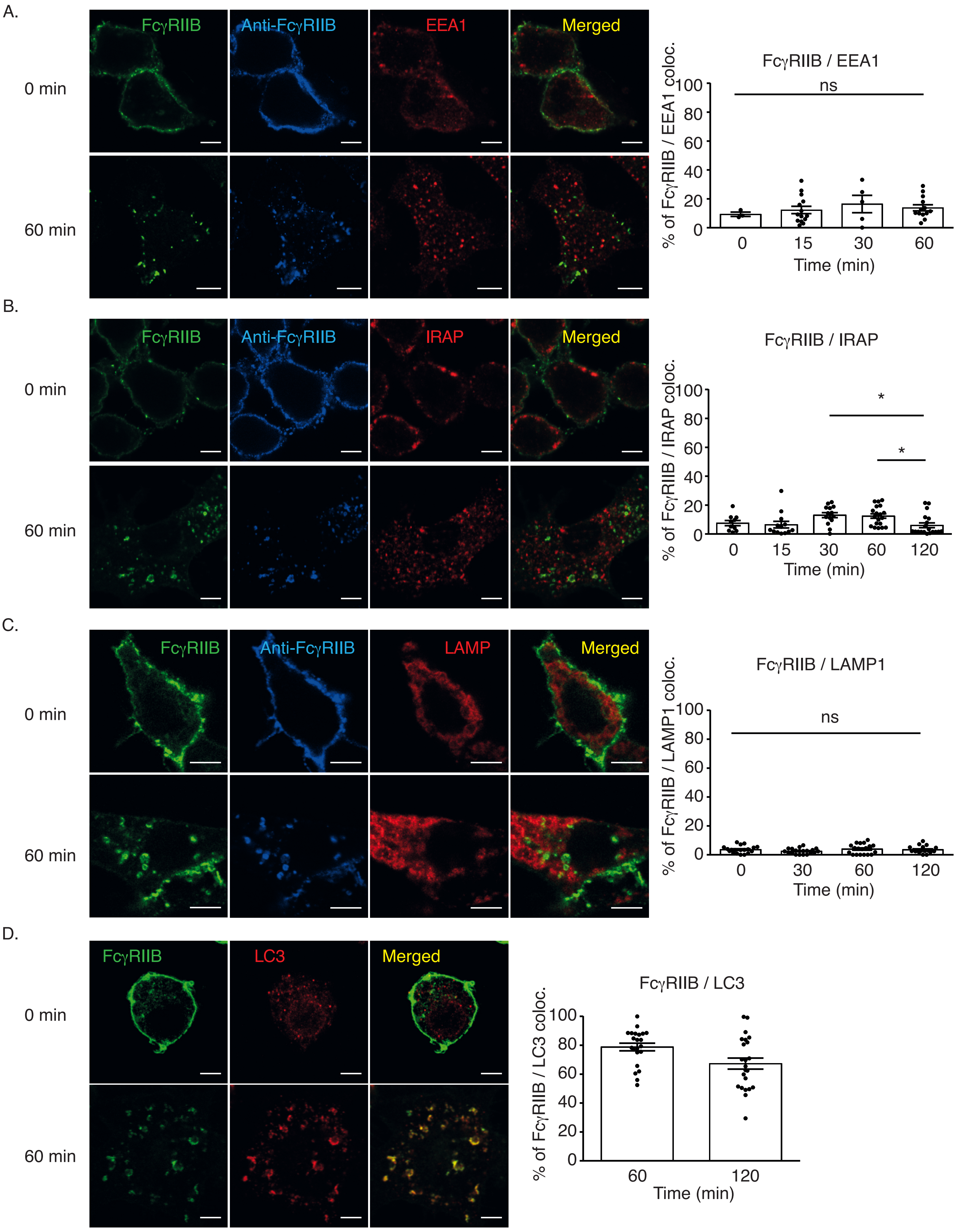


Supplementary Figure 1 Benadda et al.

### Supplementary Fig2

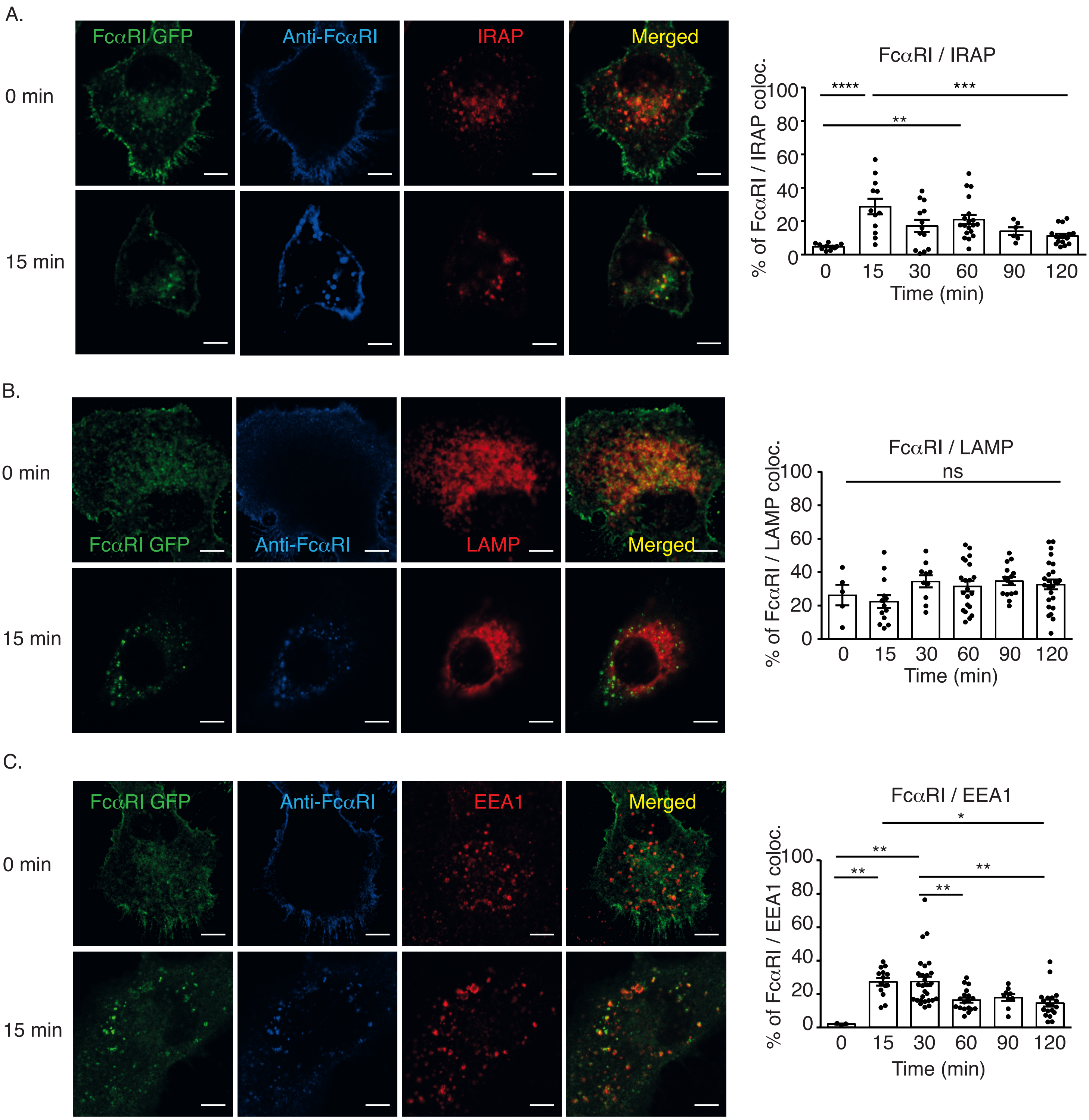


Supplementary Figure 2 Benadda et al.

### Supplementary Fig3

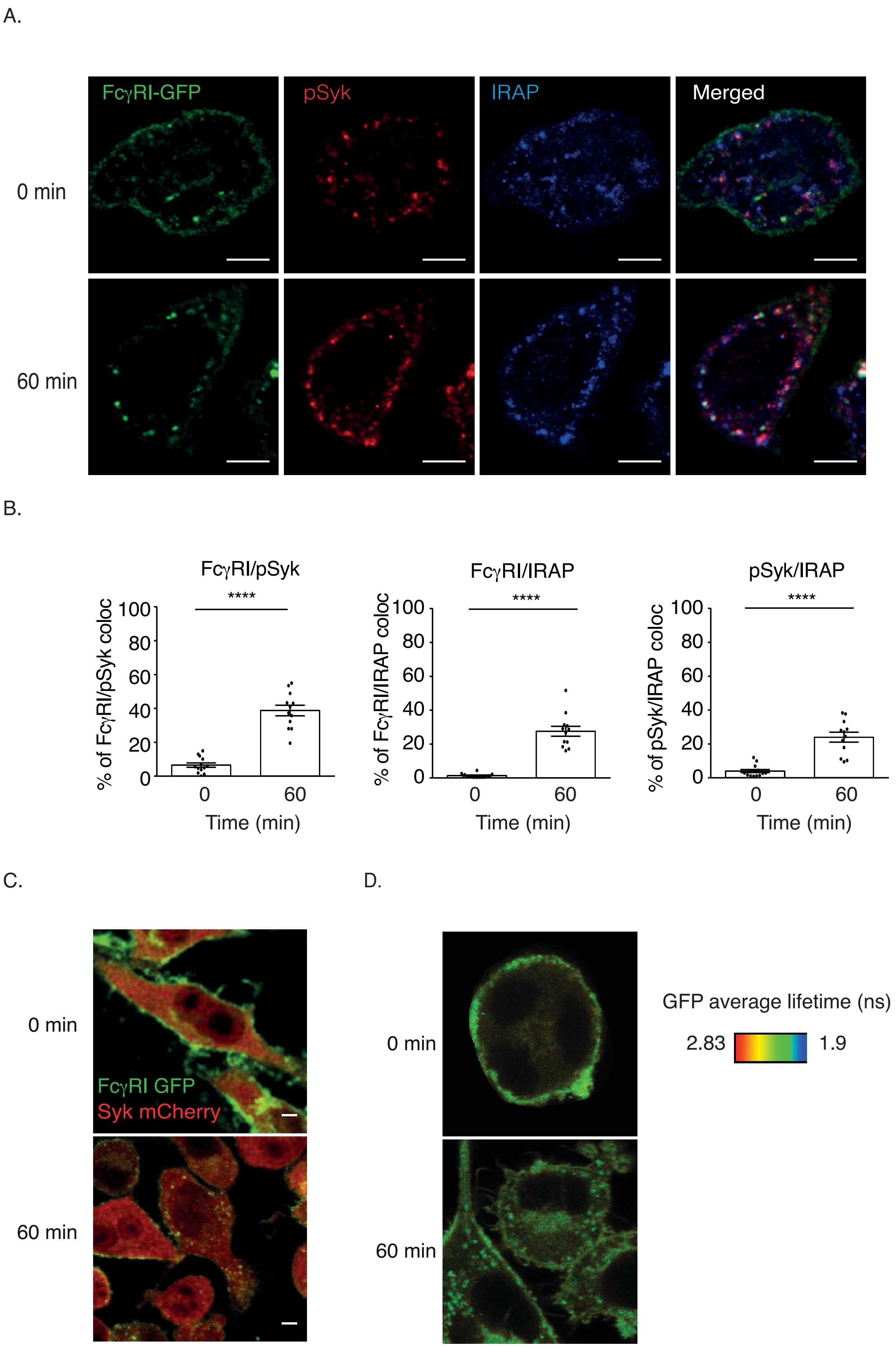


Supplementary Figure 3. Benadda et al.

### Supplementary Fig4

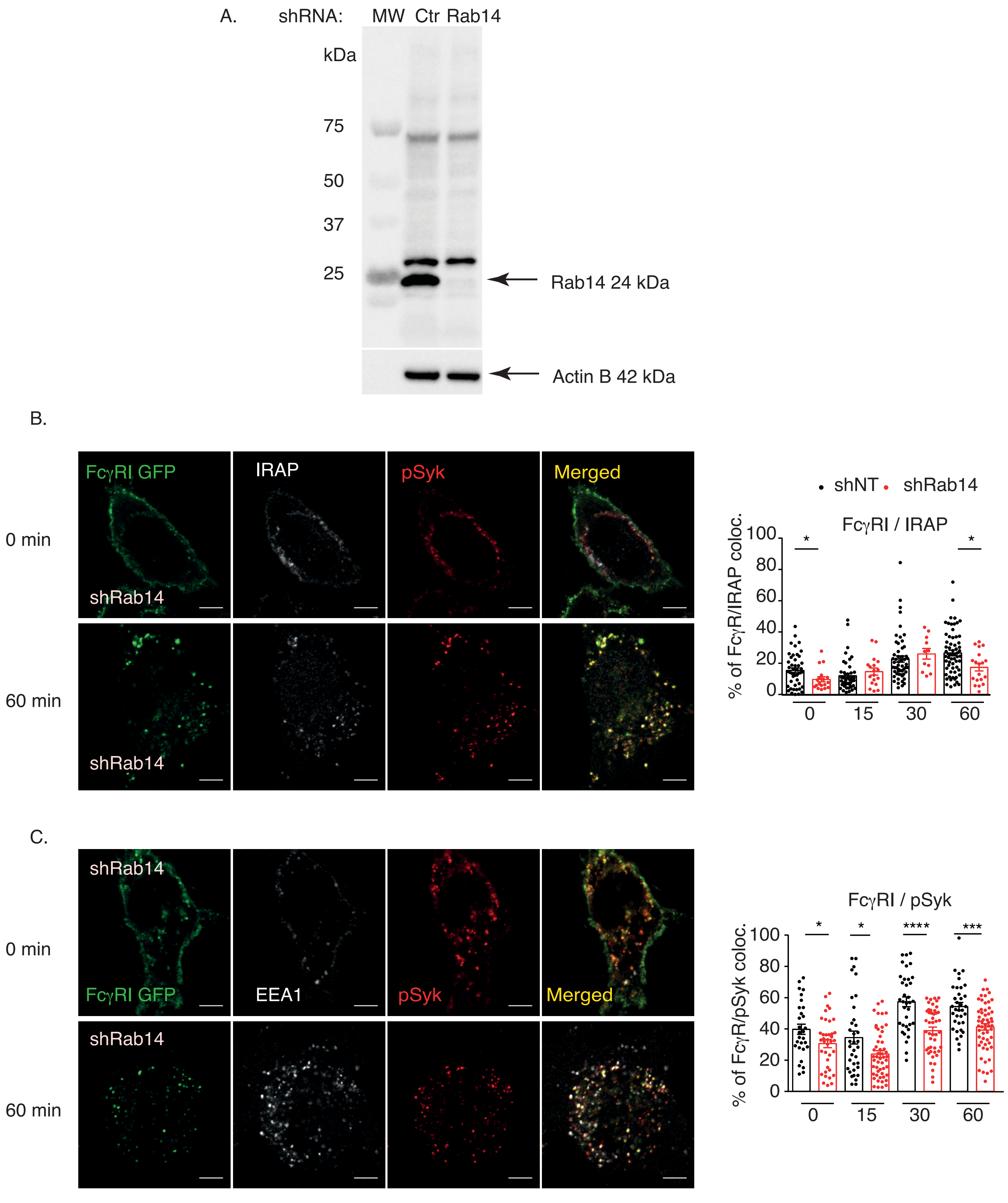


Supplementary Figure 4. Benadda et al.

### Supplementary Fig5

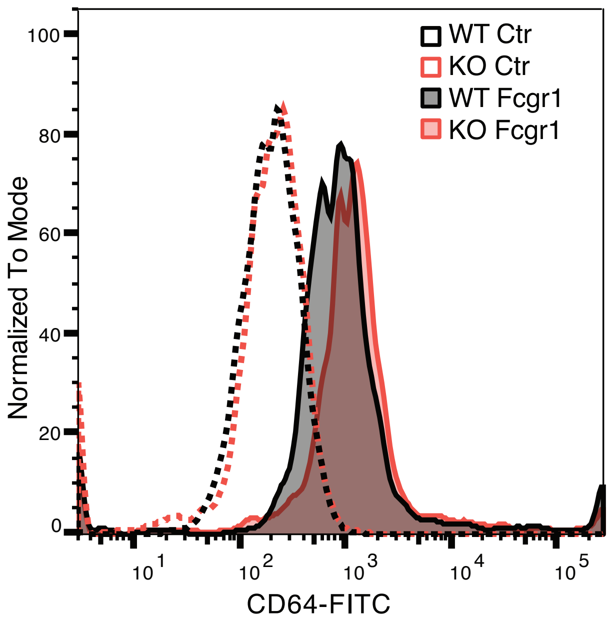


Supplementary Figure 5. Benadda et al.
